# The dynamic trophic architecture of open-ocean protist communities revealed through machine-guided metatranscriptomics

**DOI:** 10.1101/2021.01.15.426851

**Authors:** B.S. Lambert, R.D. Groussman, M.J. Schatz, S.N. Coesel, B.P. Durham, A.J. Alverson, A.E. White, E.V. Armbrust

## Abstract

Intricate networks of single-celled eukaryotes (protists) dominate carbon flow in the ocean. Their growth, demise, and interactions with other microorganisms drive the fluxes of biogeochemical elements through marine ecosystems. Mixotrophic protists are capable of both photosynthesis and ingestion of prey and are dominant components of open-ocean planktonic communities. Yet, the role of mixotrophs in elemental cycling is obscured by their capacity to act as primary producers or heterotrophic consumers depending on factors that remain largely uncharacterized. Here we introduce a machine learning model that can predict the *in situ* nutritional mode of aquatic protists based on their patterns of gene expression. This approach leverages a public collection of protist transcriptomes as a training set to identify a subset of gene families whose transcriptional profiles predict trophic status. We applied our model to nearly 100 metatranscriptomes obtained during two oceanographic cruises in the North Pacific and found community-level and population-specific evidence that abundant open-ocean mixotrophic populations shift their predominant mode of nutrient and carbon acquisition in response to natural gradients in nutrient supply and sea surface temperature. In addition, metatranscriptomic data from ship-board incubation experiments revealed that abundant mixotrophic prymnesiophytes from the oligotrophic North Pacific subtropical gyre rapidly remodelled their transcriptome to enhance photosynthesis when supplied with limiting nutrients. Coupling the technique introduced here with experiments designed to reveal the mechanisms driving mixotroph physiology is a promising approach for understanding the ecology of mixotrophic populations in the natural environment.

**Significance statement:** Mixotrophy is a ubiquitous nutritional strategy in marine ecosystems. Although our understanding of the distribution and abundance of mixotrophic plankton has improved significantly, the functional roles of mixotrophs are difficult to pinpoint, as mixotroph nutritional strategies are flexible and form a continuum between heterotrophy and phototrophy. We employ a machine learning-driven metatranscriptomic technique to assess the nutritional strategies of abundant planktonic populations *in situ* and demonstrate that mixotrophic populations play varying functional roles along physico-chemical gradients in the North Pacific Ocean, revealing a degree of physiological plasticity unique to aquatic mixotrophs. Our results highlight mechanisms that may dictate the flow of biogeochemical elements and the ecology of the North Pacific Ocean, one of the largest biogeographical provinces on Earth.

## Introduction

Single-celled eukaryotes (protists) form the base of marine food webs and their growth, demise, and interactions with other microorganisms drive the biogeochemistry of the Earth’s oceans (1, 2). Marine protists are genetically and functionally diverse, with representatives from all major lineages of the eukaryotic tree of life (3). Some protists are predatory heterotrophs that obtain organic carbon and nutrients through the ingestion of prey. They are commonly motile, consume smaller prey via phagocytic engulfment (4, 5) and may consume larger prey through veil feeding (6). Once engulfed, prey-derived nutrients are absorbed in acidic vacuoles (4, 6, 7). Other protists are strictly photosynthetic, with the ability to biosynthesize extensive suites of amino acids, lipids, pigments and other organic compounds from inorganic carbon and nutrients, with sunlight as an energy source. Photosynthesis requires a broad array of protein complexes and significant restructuring of cellular metabolism compared to that of heterotrophic organisms (8, 9), resulting in cells with distinctive cellular organization and composition. A growing list of marine protists are now described as mixotrophic and are capable of both photosynthesis and phagocytic feeding (10, 11), a trait combination rarely observed in terrestrial ecosystems (12). Marine mixotrophs are broadly distributed across the major branches of the eukaryotic tree of life, including haptophytes, dinoflagellates, chrysophytes, cryptophytes, chlorophytes, chlorarachniophytes, and ciliates (7, 10, 11, 13–17) and include many species that form harmful algal blooms (7).

Despite the pervasiveness of mixotrophy, generalizations concerning the ecology of these organisms remain elusive, given their diversity and the relative paucity of available physiological and biogeographical data. An early ‘eat-your-competitor’ hypothesis (18) suggested that by consuming bacteria, mixotrophs reduce competition for dissolved nutrients. Subsequent experimental studies augmented this concept by showing that mixotrophic species can out-compete heterotrophic specialists by grazing prey down to abundances below the critical threshold required for survival of the specialist (19, 20). Mixotrophic consumption of bacteria could benefit photosynthetic organisms as well (21), given that bacteria also compete with phototrophs for dissolved nutrients. Other laboratory-based experiments suggested that mixotrophs gain a competitive advantage over trophic specialists under nutrient-limiting conditions and sufficient light (19). Ensuing studies added additional layers of complexity by documenting species-specific differences in mixotroph grazing behaviour (22) and differential grazing responses to light and nutrient limitation (23, 24). In the natural environment, the abundance of mixotrophs is estimated through feeding experiments, where sampled communities are incubated in bottles with fluorescently labelled bacteria added as a prey source. Mixotrophs are enumerated microscopically by identifying cells that exhibit both chlorophyll autofluorescence and fluorescence due to ingested prey (11, 25). Using these approaches, mixotrophic plankton have been detected in nutrient-rich coastal waters (14, 16, 26), in coastal and open-ocean oligotrophic systems where dissolved nutrient availability is low (27, 28), and in light-limited polar waters where mixotrophy may serve as a key survival tactic for overwintering (24, 29). Within these environments, mixotrophs account for an estimated 40 to > 80% of the nanoplankton (2-20 μm) and 35 to 95% of detected bacterivory (27, 28).

The global impact of mixotrophy on carbon cycling within marine ecosystems has recently been explored through numerical models. Inclusion of organisms that can simultaneously photosynthesize and phagocytose into global ecosystem models dramatically altered modelled food web dynamics through increased efficiency of carbon transfer across trophic levels (30). This in turn led to larger average cell size in planktonic communities (30) and enhanced carbon export from surface waters (30, 31). Modelled patterns of nitrogen acquisition further suggested a shift from phagocytic acquisition of nitrogen in the North Pacific subtropical gyre to diffusive uptake northwards into nutrient rich waters. In a more explicit link between field observation and model simulations, Edwards (26) compiled data from over 100 environmental feeding assays conducted in a variety of environments and interpreted the data within a resource allocation model framework that balanced carbon and nutrient requirements under different nutritional modes. He proposed that mixotrophs outcompete obligate heterotrophs when the ratio of prey to nutrient availability is low, as mixotrophs can relieve potential carbon limitation via photosynthesis. Under conditions of nutrient limitation, mixotrophs outcompete autotrophs due to their ability to consume prey. Edwards thus predicted that mixotrophs will increase in relative abundance at low latitudes and in nutrient-rich coastal environments, while recognizing that strategies for any individual organism may vary across biogeochemical gradients. Together, these models show the potential importance of mixotrophy across a variety of environments, although the lack of high-resolution observations from diverse regions means they must rely on simplistic parametrizations of mixotrophy that are difficult to verify.

Available data emphasize the diversity, remarkable trophic flexibility, and importance of mixotrophs to marine ecosystem function. However, the diversity of mixotrophic organisms and their species-specific trophic responses to environmental conditions make it difficult to characterize the behavior of mixotrophic populations *in situ*. We approached this problem with the premise that different nutritional modes would produce distinctive transcriptional profiles. We leveraged a large public database of aquatic protist transcriptomes to develop a machine learning model that classifies trophic status based on the transcriptional profiles of key gene families. Application of this model to metatranscriptomes acquired along natural biogeochemical gradients in the North Pacific Ocean and analysis of samples from at-sea nutrient addition experiments predict that natural populations of mixotrophic protists shift their predominant mode of nutrient and carbon acquisition in response to natural gradients in nutrient supply and sea surface temperature.

## Results

### A machine learning approach to predict protistan trophic status

To determine whether transcriptional profiles could be used to distinguish between different trophic modes, we developed a transcriptome training set derived from the 678 transcriptomes of phototrophic, mixotrophic, and heterotrophic marine protists publicly available through the Marine Microbial Eukaryote Transcriptome Sequencing Project (MMETSP) (32, 33). Trophic mode labels were assigned to individual transcriptomes present in the MMETSP, allowing more than one trophic mode to be assigned to a species based on growth conditions. For example, an individual transcriptome was labelled as mixotrophic if a known mixotroph was grown in the light and in the presence of bacteria, or it was labelled phototrophic if the same species was grown in the light in the absence of bacteria. This strategy allowed us to partially decouple trophic status from phylogeny, thereby increasing the power of our model. The presumed trophic status for the majority of cells in a population at the time of sampling was assigned for ~70% of the MMETSP transcriptomes based on provided growth conditions and available literature (Table S1). The resulting training set included a total of 446 transcriptomes (275 derived from phototrophic growth conditions, 93 from mixotrophic, and 78 from heterotrophic conditions) and encompassed more than 9,000 gene families (Pfams)(34). Categorizing our training set in this manner allowed us to seek out representative features of trophic status across diverse organisms.

Gene families that significantly impacted the ability to classify trophic status were identified via the mean decrease in accuracy (MDA) algorithm (35) using two common tree-based classification algorithms: Random Forest (36) and XGBoost (37). This method of feature selection identified 901 (Random Forest) and 265 (XGBoost) candidate gene families for use in trophic classification. Based on the two classification algorithms, we defined two reduced feature sets via MDA: a common set (number of features; *f* = 120 gene families) that represented the intersection of selected gene families, and a combined set (*f* = 1046 gene families) that represented the union of selected gene families. We evaluated the ability of these two reduced sets of gene families to differentiate trophic status by applying t-distributed stochastic neighbour embedding (t-SNE) (38) to the training set. Regardless of which reduced set of gene families was examined, heterotrophic and phototrophic transcriptomes were well separated in the t-SNE latent space, with mixotrophic transcriptomes partially overlapping both clusters (Fig. S1). This separation of trophic modes was most apparent when t-SNE was applied with the common set of gene families (*f* = 120; Fig. 1A) and was not as pronounced when t-SNE was applied to transcriptomes containing all gene families (*f* = ~9000; Fig. S1D), emphasizing the importance of feature selection as a precursor to this classification task. The utility of feature selection was further highlighted when comparing the two tree-based algorithms with an Artificial Neural Network (Fig. S2, Fig. S3). Random Forest (90 ± 8%) and XGBoost (88 ± 10%) were significantly more precise than the Artificial Neural Network (77 ± 6%) (Kruskal-Wallis H test; post-hoc Wilcoxon rank sum test; *p < 0.05*), and metric values increased when reduced feature sets were used during model training (Fig. S3).

**Fig. 1.**
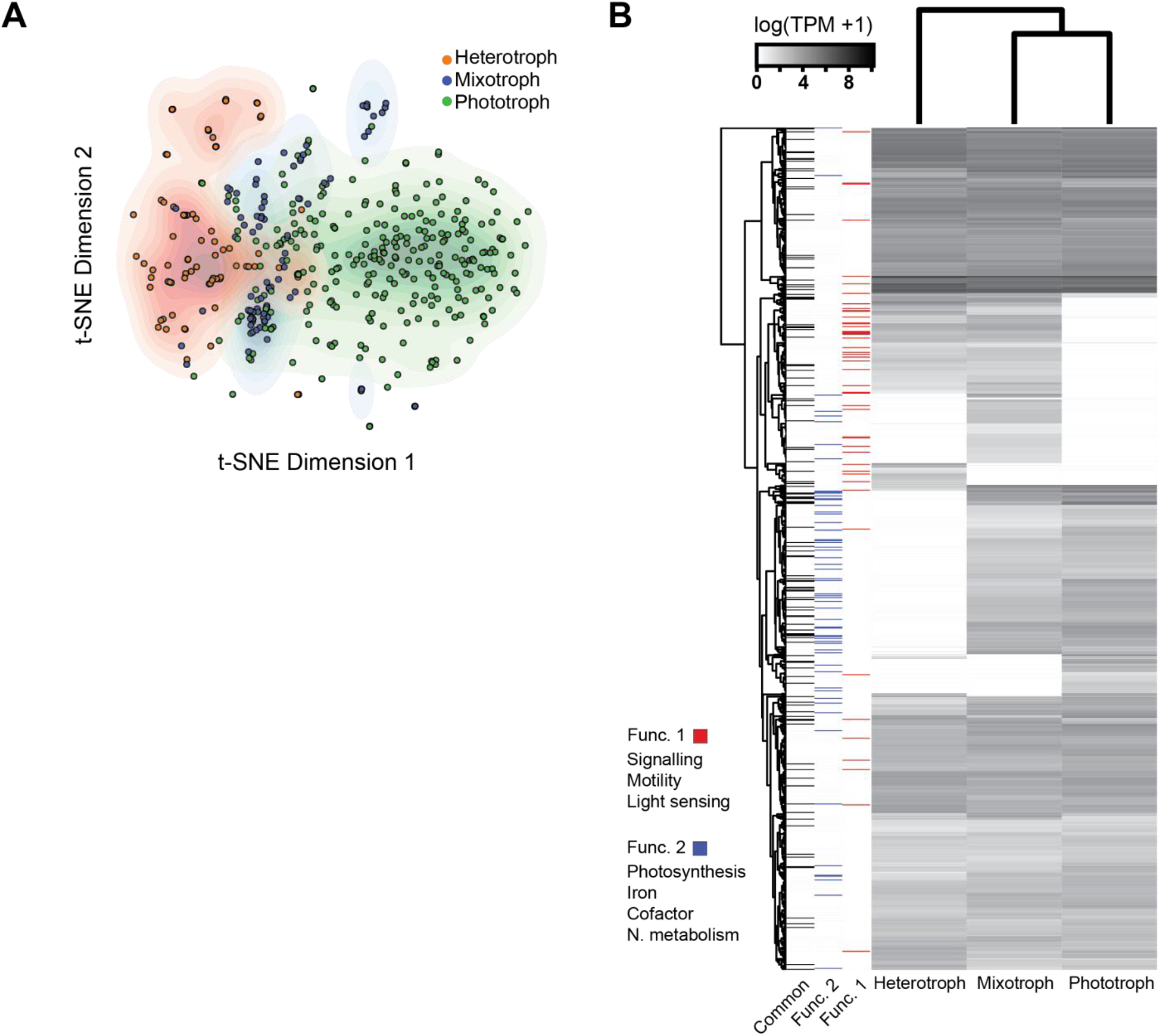
Feature selection helps distinguish trophic modes. (A) The latent representation of MMETSP phototrophic, heterotrophic and mixotrophic transcriptomes based on common selected features (*f* = 120). Transcriptional profiles of selected gene families were scaled and transformed through t-SNE. Color contours indicate kernel density for each trophic mode. (B) Median transcript abundances of the union of selected genes (*f* = 1046) in the MMETSP highlights clusters of gene families that broadly differentiate trophic modes. Gene families are annotated according to those enriched in heterotrophic/mixotrophic transcriptomes (Func. 1), enriched in phototrophic/mixotrophic transcriptomes (Func. 2), or presence in the common set of selected features. TPM: Transcripts per million.

The union of selected gene families encoded a variety of protein families, including those involved in photosynthesis, flagellar motility, and carbohydrate metabolism (Table S2). Among this reduced feature set, gene families with non-zero median expression (*f* = 886/1046) were hierarchically clustered across the MMETSP training set (Table S3) to determine whether they were differentially transcribed depending on trophic status (Fig. 1B). Relatively few gene families were enriched within heterotrophic or phototrophic specialist transcriptomes. Instead, clusters of enriched gene families with similar transcriptional profiles were shared either between phototrophs and mixotrophs or between heterotrophs and mixotrophs. Those genes families enriched in heterotrophic transcriptomes encoded proteins involved in signalling and cell cycle processes, whereas gene families enriched in phototrophic transcriptomes encoded proteins involved in lipid metabolism. Interestingly, mixotrophs displayed higher median transcription of a variety of genes that encode carbohydrate active enzymes (39), suggesting that mixotrophs may process carbohydrates in a different manner than trophic specialists. Clusters of gene families with similar relative transcript abundances within phototrophs and mixotrophs were associated with photosynthesis, cofactor synthesis, iron binding, and nitrogen metabolism, underscoring the metabolic cost of maintaining active photosystems (Fig. 1B, Func. 2). Clusters of gene families with similar relative transcript abundances in heterotrophs and mixotrophs encoded functions such as cell motility, light sensing, and signalling (Fig. 1B, Func. 1), consistent with idea that mixotrophs and heterotrophs employ motility to increase encounter with prey, rely on chemotaxis and mechanosensing to recognize prey (40, 41), and are likely attuned to the light environment to maintain exposure to light and phototrophic prey. Other selected gene families encoded proteins that were not clearly associated with a given trophic mode and included regulatory proteins, ribosomal proteins, and proteins involved in protein degradation. The common set of selected gene families encoded much of the functional diversity of the entire selected feature set (Fig. 1B; Table S3). Hierarchical clustering of the common set of gene families led to clusters that broadly shared the characteristics of clusters found in the union of selected features (Fig. S4; Table S4). The relative paucity of gene families enriched in trophic specialist transcriptomes highlights the importance of using transcriptional profiles across multiple gene families to classify trophic mode.

The performance of selected feature sets (common and union) and models (Random Forest and XGBoost) was assessed across diverse taxonomic groups and trophic modes. First, we evaluated predictions for all MMETSP transcriptomes, including the 210 MMETSP transcriptomes that could not be labelled based on available literature and thus were not part of the original training set. The two models yielded similar predictions for unlabelled transcriptomes when trained on the scaled transcript abundances of either the union or common set of selected features (Cohen’s κ = 0.90, 0.93 respectively; Table S5) (42), which was a significant improvement over training with all ~9000 gene families (Cohen’s κ = 0.51). To determine how trophic mode predictions were distributed across taxa, an 18S rDNA phylogenetic tree was constructed for each transcriptome present in the MMETSP. When available, leaves were labelled by the assigned trophic mode based on our literature review (training label; Table S1), as well as the predicted nutritional mode when predictions from both Random Forest and XGBoost models trained on the union of selected features agreed. As expected, the model-predicted trophic mode was in accordance with the literature-established trophic mode for taxa used in the training set (Fig. 2A). For the unlabelled transcriptomes, the greatest consistency between the two algorithms was found within phyla dominated by either phototrophs or heterotrophs. Classifications were inconsistent between the two models for a subset (10%) of transcriptomes derived primarily from species distributed across the dinoflagellates and the haptophytes, two phyla that include known phototrophs, mixotrophs, and heterotrophs (10).

**Fig. 2.**
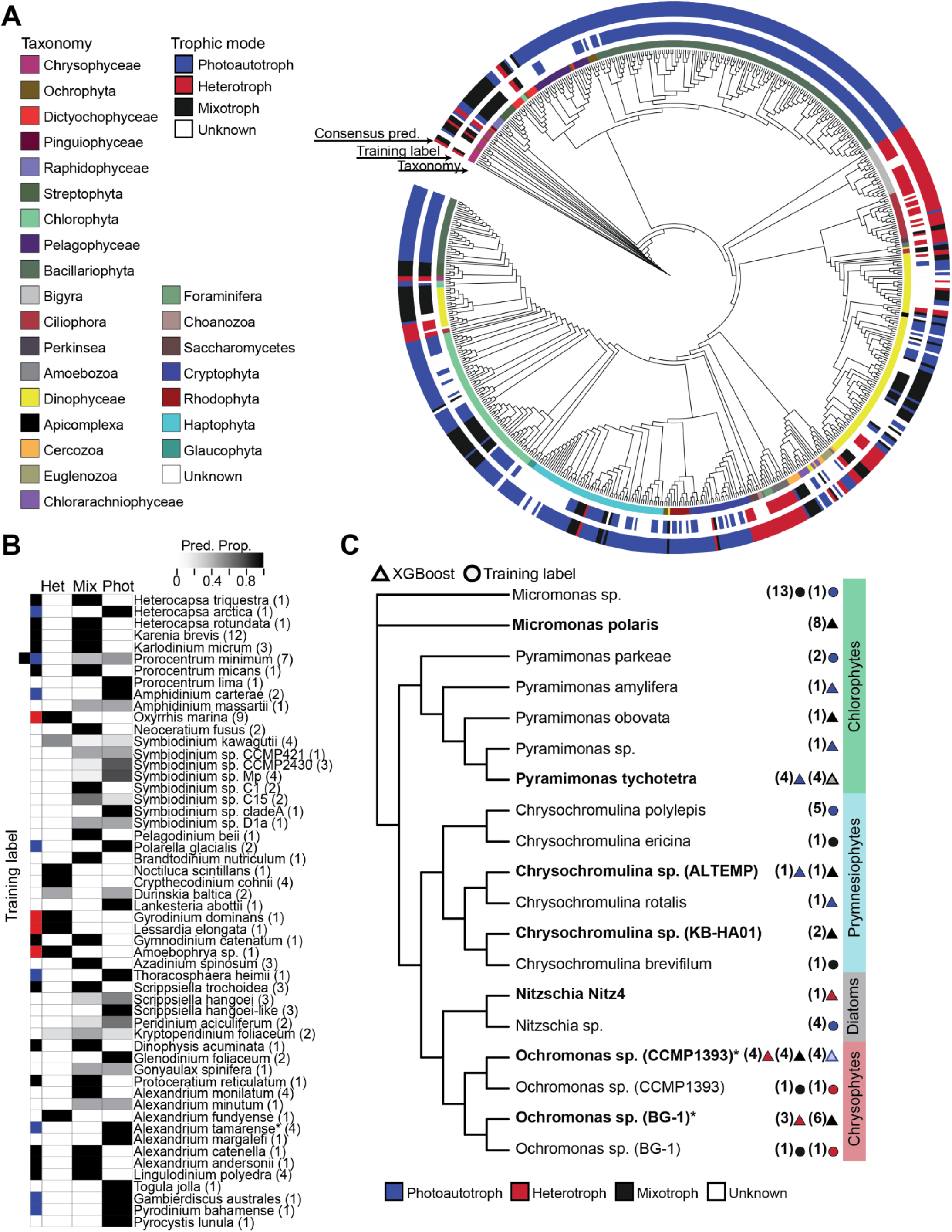
Trophic mode predictions for MMETSP transcriptomes and culture-derived validation transcriptomes. (A) Phylogenetic representation of all taxa represented in the MMETSP database. Color strips surrounding the tree indicate (from inner to outer): taxonomic group membership, transcriptome trophic mode training label (if present), and consensus trophic mode prediction between RF and XGBoost (if present) using the union of selected gene families. (B) Dinoflagellate taxa represented in the MMETSP database. Indicated are the training labels, when available (annotation strip; blue: phototrophy; red: heterotrophy; black: mixotrophy) and the proportion of trophic mode predictions derived from XGBoost model output using both selected feature sets. \**A. tamarense* was only grown axenically. *P. minimum* transcriptomes were derived from both phototrophic and mixotrophic culture conditions. The number of transcriptome replicates available in the MMETSP are indicated in parentheses. (C) Phylogenetic representation of validation taxa (bold) known to exhibit diverse trophic modes and their most closely related MMETSP representatives. The validation species *Nitzschia* sp. Nitz4 represents a non-photosynthetic diatom. The chrysophyte *Ochromonas* sp. (strains BG-1 and CCMP1393) and chlorophytes *Micromonas polaris* and *Pyramimonas tychotetra* have the capacity for mixotrophy. Values in parentheses indicate the number of transcriptomes present in the MMETSP. Colored circles to the right indicate trophic mode labels for MMETSP organisms in the training set. Triangles indicate XGBoost predictions for unlabelled transcriptomes. Translucent triangles indicate that the condition was present, but XGBoost made an incorrect prediction. *Species represented in the MMETSP.

We therefore further inspected predictions for the MMETSP dinoflagellate transcriptomes (XGBoost: Fig. 2B; Random Forest: Fig. S5), which were derived from organisms primarily grown in the light in the presence of bacteria (Table S1). About 75% of the 56 unlabelled dinoflagellate transcriptomes were consistently classified by the two models as either phototrophic (16), heterotrophic (9) or mixotrophic (17). Organisms with mixed classifications (derived from XGBoost and both reduced feature sets; Table S5) predominantly received a combination of mixotrophic and heterotrophic classifications or mixotrophic and phototrophic classifications. These two types of mixed classifications were most apparent within the unlabelled species of the genus *Symbiodinium*, which includes both free-living species and those living in symbiosis with diverse hosts. Transcriptomes from *S. kawgutii* were primarily classified as heterotrophic (~ 63%), with a small proportion of transcriptomes classified as either phototrophic or mixotrophic. Transcriptomes from all other *Symbiodinium* species were classified as predominantly phototrophic or predominantly mixotrophic, consistent with a phototrophic lifestyle that includes the potential for induced phagotrophy (29, 43). Similarly, two members of the *Alexandrium* species complex had predictions split between phototrophy and mixotrophy, which agrees well with observations of both trophic modes within this group (7, 44), whereas *Alexandrium fundyense* (*n* = 1) was predicted to be a heterotroph, a result inconsistent with the literature. Both models predicted *Neoceratium fusus* to be mixotrophic, a strategy observed for this species in polar waters (24). Two species received classifications that were in logical conflict: *Kryptoperidinium foliaceum* (*n* = 2) received model predictions split across all trophic modes and *Durinskia baltica* (*n* = 2) received classifications split between heterotrophy and phototrophy. Interestingly, both dinoflagellates are known ‘dinotoms’ that retain the nucleus, plastid, and mitochrondria of a diatom endosymbiont (45), perhaps contributing to the mixed predictions. Predictions split between phototrophy and heterotrophy or across all trophic modes are difficult to interpret and could reflect complicated evolutionary histories such as tertiary symbioses, true mixotrophy, trophic heterogeneity, or transcriptional profiles that conflict with the decision boundaries of our models. Therefore, to be conservative with predictions, we did not attempt to classify trophic status when replicate transcriptomes resulted in classifications split between phototrophy and heterotrophy. This result highlighted the importance of transcriptome replicates to build classification confidence based on prediction consistency. Our final test with the MMETSP transcriptomes was to evaluate whether the models could predict osmoheterotrophy, a process by which heterotrophic organisms sustain growth through the uptake of dissolved compounds rather than by consumption of prey. MMETSP osmoheterotrophs (non-photosynthetic organisms that acquire carbon via diffusive uptake) were purposely withheld from the training set (Table S1) and used as a test of model generalization. The transcriptomes of these osmoheterotrophic organisms were consistently predicted to be heterotrophs, despite the absence of osmoheterotroph transcriptomes in the training data, confirming that both models generalized well in making predictions of heterotrophy, despite relatively few examples in the training set.

We next challenged the models with transcriptomes not present in the MMETSP dataset (Fig. 2C). Because all diatoms within the MMETSP are obligate phototrophs, we tested the models with a transcriptome from a known non-photosynthetic diatom *Nitzschia* sp. (Nitz4; Fig. S6) (46). Both Random Forest and XGBoost successfully classified this diatom as heterotrophic despite the presence of four photosynthetic *Nitzschia* species within the training set, reiterating the ability of the model to disentangle phylogeny and trophic status. Transcriptomes from *Ochromonas* sp. (strains BG-1 and CCMP1393), *Micromonas polaris*, and *Pyramimonas tychotetra* were chosen as they are known mixotrophs, and in each instance, experiments were carried out that modulated the availability of light, prey, or nutrients in an effort to drive organisms into distinct nutritional modes (Table S6) (22, 47, 48). Particle ingestion by *M. polaris* was detected in both the high- and low-nutrient conditions employed (48) and both models accordingly classified *M. polaris* from both conditions as mixotrophic. *Pyramimonas tychotetra* (Fig. S6) was cultured under high- or low-nutrient conditions, with the low nutrient condition intended to induce phagotrophy though only low levels of bacterivory were observed (48). Both models accurately predicted phototrophy in the high nutrient condition. However, both models also predicted phototrophy in the low-nutrient condition, which could reflect either the limited bacterial grazing documented in the experiment (48) or the absence of representative mixotrophic *Pyramimonas* sp. transcriptomes in the training set. Two *Ochromonas* species were also grown under different conditions expected to select for either autotrophic, heterotrophic, or mixotrophic growth. Both models predicted the expected nutritional mode for all conditions except when *Ochromonas* strain CCMP1393 was grown in the absence of bacteria, which was expected to induce pure autotrophy. Lastly, we recognized that the MMETSP data set was derived primarily from coastal organisms. To evaluate the applicability of these models to open ocean species, we assembled, annotated, and classified transcriptomes from two *Chrysochromulina* sp. isolated from coastal Hawaii (KB-HA01) and the North Pacific subtropical gyre (AL-TEMP). One isolate was predicted to be mixotrophic (KB-HA01) and the other had predictions split between phototrophy and mixotrophy (AL-TEMP), consistent with observed constitutive mixotrophy in a variety of *Chrysochromulina* species and data from our laboratory (Fig. S7). XGBoost consistently outperformed Random Forest when applied to validation transcriptomes (81% vs 74% accuracy) and thus was selected for subsequent analyses together with the union of selected gene families (*f* = 1046).

Collectively, these tests identified several elements that impact model performance. First, accurate classification required transcriptional profiles of numerous selected gene families, many of which have either no known function or a predicted function with no obvious relation to nutritional status, and transcriptional patterns of individual genes were not sufficient for classification. The requirement for a holistic approach to trophic mode prediction is highlighted by comparison to a recent study that proposed 4 genes required for phagocytosis during bacterivorous feeding by heterotrophic organisms (49). Two of the four proposed phagocytosis marker gene families (Pfam IDs: PF09286, PF03030) were part of our selected feature set, and the transcriptional profile of only one of these genes was specific to mixotrophy and heterotrophy. Second, prediction consistency between models and prediction accuracy increased when predictions were made for organisms closely related to those present in the MMETSP training set. Third, greater confidence in assigning trophic status was achieved through increased numbers of transcriptome replicates, often by yielding a clear majority vote prediction. Last, in those instances where replicate transcriptomes resulted in predictions that included both phototrophy and heterotrophy, we assumed that these transcriptional profiles were in conflict with the decision boundaries of our model and that nutritional status could not be predicted accurately for these transcriptomes.

### Predicting the in situ nutritional status of marine planktonic populations

The ability of our model to predict the trophic status of model organisms based on validation transcriptomes obtained from differing growth conditions provided both confidence and an understanding of the constraints for applying this approach to transcriptomes derived from natural communities. Metatranscriptomes provide a snapshot of community function and metabolism and can be deconvolved into transcriptomes associated with representative taxonomic bins. A total of 95 eukaryotic (polyA-selected) metatranscriptomes were collected on two field campaigns in the North Pacific Ocean. The metatranscriptomes were deconvolved into transcriptomes from 26 species-level taxonomic bins based on Lowest Common Ancestor assignment (Fig. S8). Further analysis was conducted with those taxonomic bins present at any given sampling site with a minimum of 4 transcriptomes present in the dataset that met established completeness criteria (Note S1) and were closely related to transcriptomes present in the original training set. The union of selected features (*f* = 1046) was used for classification.

Samples for 48 of the metatranscriptomes were collected during the SCOPE Diel Cruise (KM1513; HOELegacy 2, July/Aug, 2015; Fig. S9), from near Station ALOHA (158 °W, 22.45 °N) in the North Pacific subtropical gyre, a low latitude oligotrophic region where the availability of nitrogen limits productivity in surface waters and where ecosystem models (30) and observational studies (25, 27, 50) indicate the potential for abundant and active mixtotrophic organisms. Duplicate samples (0.2 – 100 μm) were collected from a depth of 15 meters every 4 hours over 4 days following a Lagrangian drifter. Eighteen species-level taxonomic bins (Table S7; Fig. S10) were retrieved from these metatranscriptomes (SI Methods) that met our established completeness criteria (Note S1), including detection of transcripts associated with the selected feature set (1023 of the 1046 features were detected in transcriptomes belonging to the extracted taxonomic bins). Most (14) of the taxonomic bins were present throughout the day-night cycle of the 4-day sampling period and were each represented by at least 32 transcriptomes. A total of 13 taxonomic bins corresponded to reference species with transcriptomes labelled as mixotrophic in the training set or else predicted as mixotrophic in the MMETSP dataset. The remaining 5 taxonomic bins corresponded to reference species with transcriptomes designated as heterotrophic (1) or phototrophic (4) based either on training labels or predictions from the MMETSP – although it should be noted that transcriptomes from 3 of the reference species classified as phototrophic in the training dataset were mixotrophic species grown in the absence of bacteria. Thus, most of these environmental species had the potential to exhibit broad-ranging nutritional strategies.

Nine environmental dinoflagellate bins were predicted to behave primarily either as heterotrophs (6 species) or as phototrophs (3 species), regardless of whether the transcriptomes were derived from day or night samples (Fig. 3A, Table S8), suggesting that these dinoflagellates were behaving as trophic specialists at the time of sampling despite their presumed capacity for mixotrophy. A similar phenomenon was observed with the five environmental prymnesiophyte bins, four of which also have the capacity for mixotrophy based on laboratory studies (50, 51). *Chrysochromulina brevifilum* was the only haptophyte species predominantly classified as a mixotroph at the time of sampling, all other prymnesiophytes were predicted to have behaved primarily as heterotrophs (2) or as phototrophs (2) when sampled. Transcriptomes from two environmental dinoflagellate bins – the mixotrophic *Scrippsiella trochoidea* and the heterotrophic *Oxyrrhis marina* – resulted in conflicting predictions of phototrophy and heterotrophy (> 25% for both categories across all classified transcriptional profiles), while lacking mixotrophic predictions. This suggested that the transcriptional profiles for these two taxonomic bins lie close to model decision boundaries (Note S1), and according to defined criteria, were not considered further (~12% of the total retrieved transcriptomes; see Fig. S11 for profiles including these bins). Finally, dictyophyte field predictions were consistent with MMETSP-derived labels: *Rhizochromulina marina* transcriptomes were classified as mixotrophic and *Dictyocha speculum* as a mixture of phototrophic and mixotrophic. The stability of predictions over day/night cycles despite known diel oscillations in transcript abundances (51) further indicated that our model leverages a wide array of transcriptional features when predicting nutritional mode. Overall, predictions of heterotrophy dominated when all predictions were aggregated over each individual 24 hr period (*n* = 4) and prediction proportions remained stable over the diel period (Fig. 3B). Thus, this low-nutrient environment with abundant sunlight appeared to select for a predominance of heterotrophic feeding with less reliance on photosynthesis or mixotrophy, consistent with the hypothesis that phagotrophy provides protists with necessary carbon/nutrients in oligotrophic environments.

**Fig. 3.**
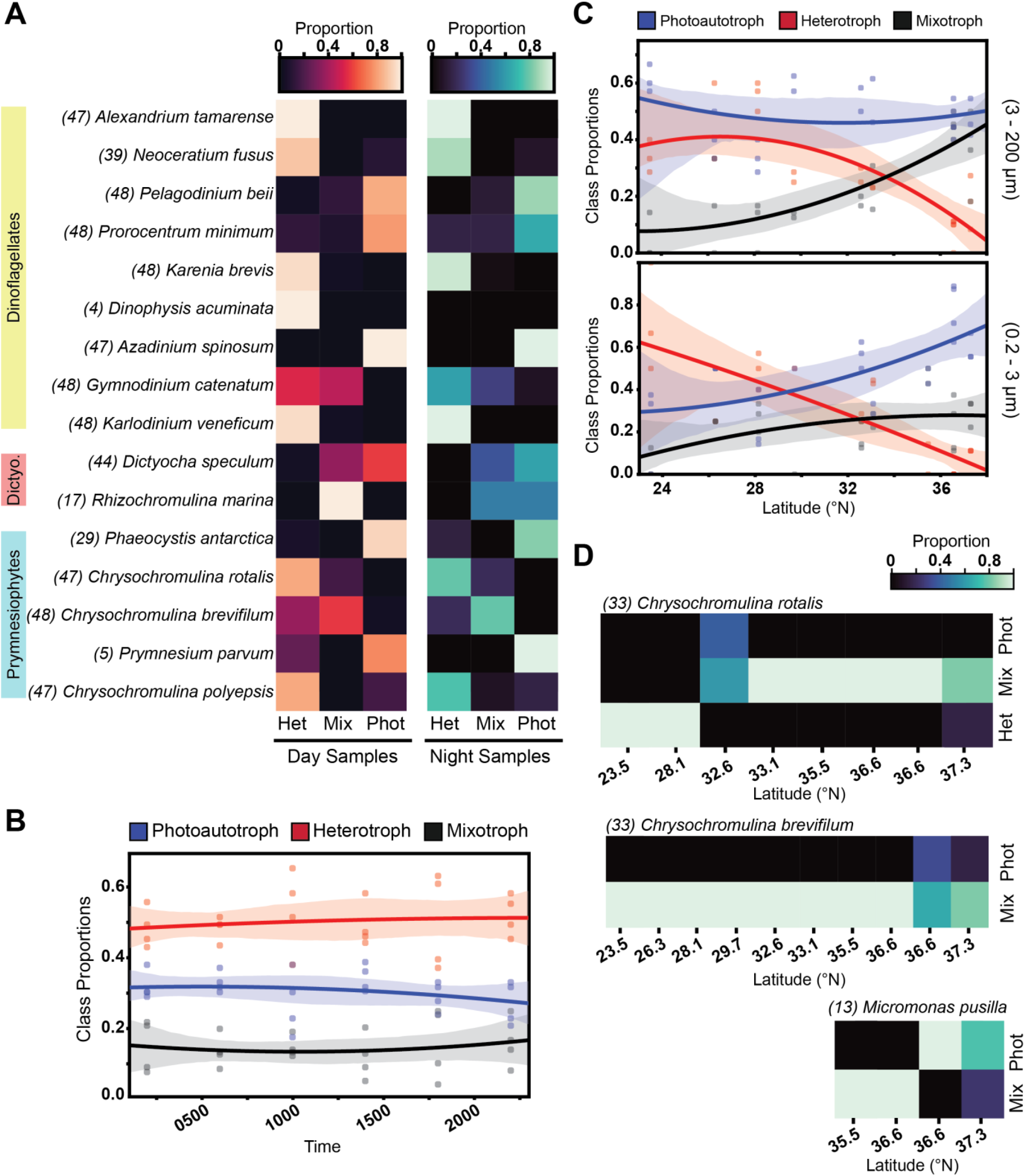
Distribution of model predictions across environmental taxonomic bins. (A) Diel transcriptomes (KM1513, July/Aug. 2015, 22 °N) were binned according to period of sample acquisition. Heatmaps show the proportion of XGBoost-predicted trophic modes for each transcriptome bin for samples taken during the day (left) vs night (right). The number of transcriptional profiles classified per taxonomic bin is indicated in parentheses. (B) Proportion of trophic mode predictions (derived from (A)) over a diel cycle. Individual points represent samples obtained at the same time of day (*n* = 4). (C) Gradients 1 predictions were summed by latitude and normalized by total number of predictions at each site to generate trophic mode prediction proportions. Second order regressions are represented by solid lines and individual station values are plotted as points; 95% confidence intervals are displayed as shaded regions surrounding fitted curves. (D) Proportion of trophic mode predictions for three abundant protists across the G1 transect. The number of transcriptional profiles classified per taxonomic bin is indicated in parentheses.

To test whether species and community-level trophic status changed under varying environmental conditions, we used SCOPE Gradients 1 (KOK1606, April/May, 2016; Fig. S9) metatranscriptome samples collected at 15 meters depth from 10 stations (see Table S9 for replication scheme) spanning from 23.5 °N to 37.3 °N along 158 °W, traversing from the oligotrophic subtropical gyre to the more nutrient-rich region of the North Pacific transition zone (Fig. S9). The southernmost stations of Gradients 1 are within the gyre, ~ one degree north of Station ALOHA, with similar community composition to that observed on the Diel cruise. A total of 47 metatranscriptomes (Table S9) were deconvolved into 437 transcriptional profiles representing 23 species-level taxonomic bins (Table S7; Fig. S10) distributed between small (0.2 – 3 μm) and large (3 – 200 μm) size class samples (47% vs 53% respectively), with ~98% of the selected features present across the collection of retrieved profiles. We examined environmental taxonomic bins at each station along the Gradients 1 transect, with the same completeness criteria and exclusion criteria of conflicting phototrophy/heterotrophy classifications. To filter incongruous predictions, predictions for each taxonomic bin were grouped by latitude. When both heterotrophic and phototrophic predictions were present for a taxonomic bin in a proportion > 25% for a given latitude, all transcriptional profiles for the taxonomic bin were removed at that latitude. In this manner, a total of 43 transcriptomes belonging to 7 taxonomic bins were excluded from the large size fraction and 23 transcriptomes belonging to 4 taxonomic bins were excluded from the small size fraction (removing a combined ~15% of the retrieved transcriptomes). For both cruises, exclusion of these bins did not affect community-level patterns (Fig. S11). Of the 23 recovered taxonomic bins, 15 were also present in the Diel dataset.

In both size fractions, a significant reduction in the proportion of heterotrophic predictions was observed with increasing latitude (*p* ≪ 0.01; Fig. 3C). In the large size fraction, this was predominantly accompanied by an increase in predictions of mixotrophy. In the small size class, both mixotrophy and phototrophy predictions increased northwards (Table S10). These changes could reflect differences in community composition, shifts in widespread populations’ nutritional strategies, or both. We therefore evaluated the *in situ* nutritional modes of three taxonomic species bins – *Chrysochromulina rotalis, Chrysochromulina brevifilum*, and *Micromonas pusilla* – that were broadly distributed across sites. Within the gyre (demarcated at ~30.5° N by the salinity front; Fig. S9), the two *Chrysochromulina* species bins displayed contrasting strategies, with *C. rotalis* predicted to be primarily heterotrophic and *C. brevifilum* predicted to be mixotrophic (Fig. 3D), in agreement with the predictions for these same environmental species sampled during the Diel cruise in the summer of the previous year (Fig. 3A). Within the more nutrient-rich waters of the transition zone, we observed a shift to a greater reliance on photosynthesis, reflected in mixotrophy predictions for *C. rotalis* and increasingly split predictions between phototrophy and mixotrophy for *C. brevifilum* at higher latitudes. Although the latitudinal range of *M. pusilla* was not as broad, we also observed a distinct shift in nutritional mode predictions from mixotrophy to phototrophy at higher latitudes. These results supported the hypothesis that nutrient availability is a key driver of the nutritional mode employed by these flexible organisms in the surface waters of the North Pacific.

### Environmental drivers of planktonic nutritional strategies

Multiple environmental parameters are correlated with a transition from the North Pacific oligotrophic gyre to the more nutrient-rich waters of higher latitudes. The Gradients 1 cruise spanned gradients in inorganic nutrient concentrations, microbial biomass, surface ocean temperature, and surface light levels (photosynthetically available radiation or PAR). Available nitrate at the time of sampling increased north of the subtropical gyre with a counter-gradient in iron (Fig. 4A; PISCESv2 output (52) and measured), which increased southwards into the subtropical gyre. Both photosynthetic (satellite-derived) and bacterial (measured) biomass increased northwards into the more productive transition zone waters (Fig. 4B), coincident with a sharp increase in *Synechococcus* abundance at the gyre boundary (Fig. 4C). Changes in biomass composition in response to increased nutrient availability were indicated by particulate carbon:nitrogen (C:N) and carbon:phosphorus (C:P) that were greater than the Redfield ratio of 106 C:16 N:1 P (53) within the subtropical gyre (54) (~30.5 °N) and decreased to below (C:N) or to (C:P) the Redfield ratio within the transition zone (Fig. 4D). The northwards increase in biomass and productivity occurred against a backdrop of decreasing sea surface temperature and light availability (Fig. 4E; Satellite-derived).

**Fig. 4.**
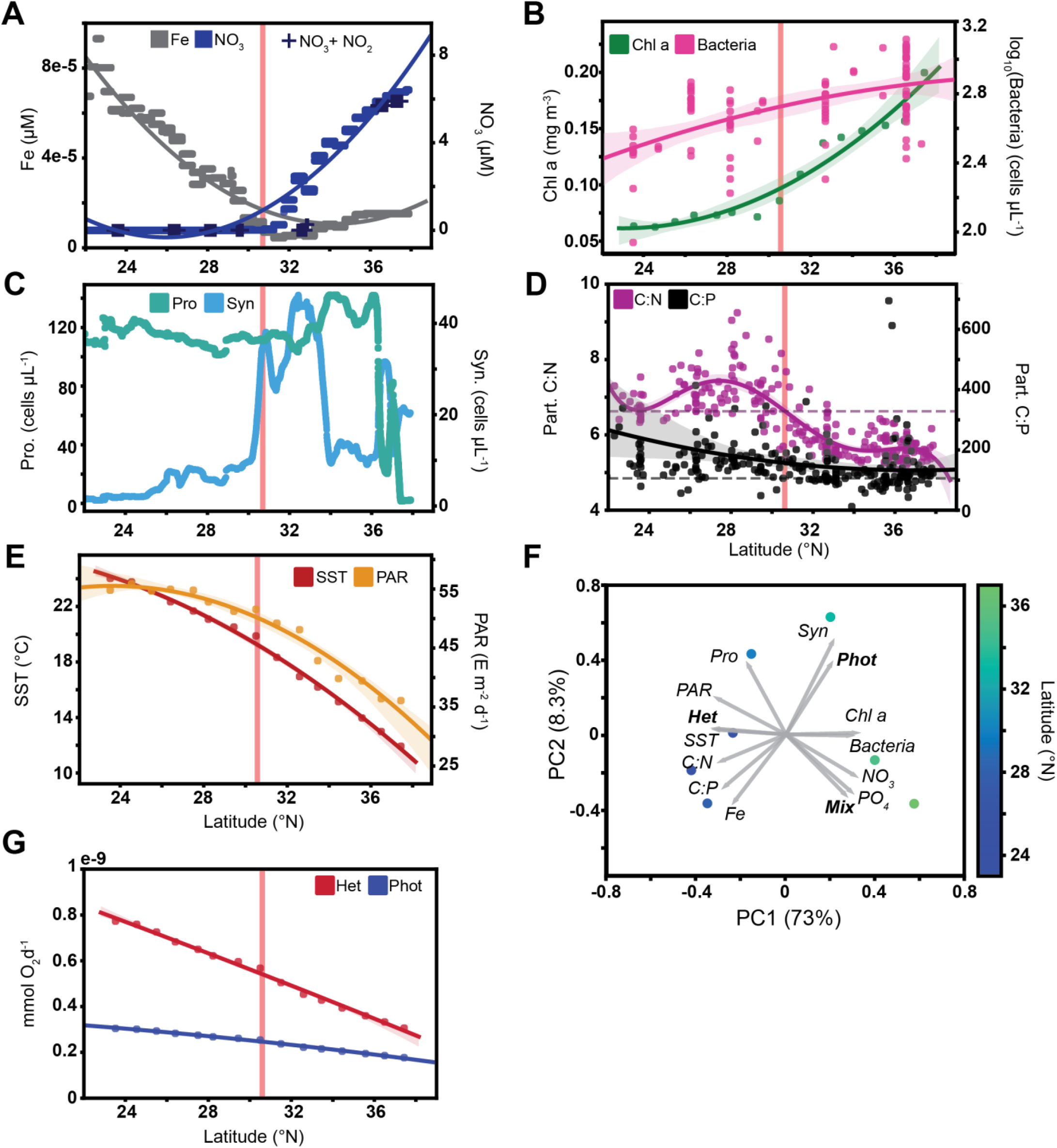
Environmental drivers of trophic mode in Gradients 1 surface waters. (A) Counter-gradients in Fe and NO_3_ across latitude as demonstrated by modelled data (line) and observations (+). Model data was obtained from the PISCES-v2 biogeochemical model and experimental nitrate + nitrite was determined through a chemiluminescence method (85) for samples obtained during the Gradients 1 cruise. Solid lines represent a 2^nd^ order fit to the data and the shaded region indicates the 95% confidence interval. (B) Photosynthetic biomass proxy *chl a* concentration calculated from MODIS satellite data products (SI Methods) and heterotrophic bacterial counts determined *via* flow cytometric analyses of seawater samples. Solid lines represent a 2^nd^ order fit to the data and the shaded region the 95% confidence interval. (C) *Synechococcus* abundance and *Prochlorococcus* abundance derived from the shipboard flow cytometer SeaFlow. (D) Particulate C:N and C:P. Transition in particulate C:N from greater than the Redfield ratio (indicated by dashed purple line) to below coincides with the salinity front near 30°N (red vertical line in each plot). Solid lines represent a 5^th^ order fit (C:N) and a 2^nd^ order fit (C:P) to the data with shaded regions representing the 95% confidence intervals. (E) PAR and sea-surface temperature (SST) derived from satellite products (77, 78). Solid lines represent a 2^nd^ order fit to the data and the shaded region the 95% confidence interval. (F) PCA bi-plot showing the relation between trophic mode predictions and environmental metadata. Samples were localized to the nearest degree latitude and averaged. Remaining stations with data available across all datasets are plotted as points in the latent space, while environmental parameters and trophic mode proportions are represented as overlaid vectors. Stations are colored by latitude. (G) Derived metabolic scaling relationships predict a disproportionate temperature-driven decrease in respiration rates for heterotrophs with increasing latitude. Solid lines are a linear fit of the data and the shaded regions indicate the 95% confidence interval. See Note S2 for more details concerning metabolic rate calculations.

To identify potential drivers of trophic mode, a principal component analysis was conducted by incorporating trophic status predictions with measured and modelled environmental parameters (SI Methods; Fig. 4F). The proportion of heterotrophic predictions strongly aligned with the first principal component (explaining 73% of the variance), alongside sea surface temperature and other indicators of subtropical conditions. This result is consistent with the Metabolic Theory of Ecology (MTE), which predicts a disproportionate decrease in respiration rates for heterotrophic organisms with decreasing temperature (Fig. 4G; Note S2), suggesting a reduction in heterotrophic respiration into the cooler waters of the North. Indeed, a disproportionate reduction in respiration rates for heterotrophic protists is expected for the light and temperature fields associated with the higher latitudes sampled along the Gradients 1 transect (Fig. 4G). Phototrophic predictions primarily coincided with increasing *Synechococcus* abundance, consistent with the transition out of the subtropical gyre, but prior to more northward increases in available nitrate and phosphate concentrations. Mixotrophy predictions aligned with increasing dissolved nitrate and phosphate concentrations (derived via the PISCESv2 biogeochemical model) (52) and decreasing PAR, a result that runs counter to an earlier assumption that mixotrophy would be most competitive in low-nutrient, high-light waters (19). We propose that these results instead reflect a combination of temperature dependencies and the quality of prey items relative to dissolved nutrient availability. Optimal Foraging Theory (OFT) (55) predicts that in patchy resource environments, prey quality (*i.e*. particulate C:N for nitrogen-limited phagotrophs) influences whether mixotrophic organisms rely primarily on nutrients derived from prey via phagotrophy or on dissolved nutrients and photosynthesis (56). Together, our results suggest that nitrogen limitation in the subtropical gyre drives mixotrophic organisms to rely primarily on phagotrophy to meet nitrogen and organic carbon demands, consistent with the dominance of heterotrophic predictions in these waters. At higher latitudes, increased nutrient availability supports photosynthesis by mixotrophic organisms, potentially offsetting the increased costs associated with continued phagotrophy as temperature decreases.

Trophic mode predictions for environmental taxonomic bins suggested that an increase in dissolved nutrient availability would cause mixotrophic organisms to shift their dominant trophic mode. We tested this hypothesis using metatranscriptomes derived from on-deck incubation experiments carried out on the SCOPE Gradients 2 cruise (MGL1704; May/June, 2017; Fig. S9) in which the nitrogen-limited subtropical gyre community was supplemented with nitrogen and phosphorous. Ship-board incubations carried out with the gyre’s microbial community allowed us to decouple shifts in nutrient availability from both community compositional changes and shifts in environmental covariates along the cruise transect. Samples were collected before (t = 0) and after (t = 96 hr) nutrient amendments. Logistical constraints (*i.e*. reduced sequencing depth, low biomass) and smaller sample sizes meant that the resulting metatranscriptomic data allowed for analyses of several mixotrophic prymnesiophytes binned together rather than as individual species. Amendment of samples with 5 μM nitrate and 0.5 μM phosphate led to significant (*p_adj_* < 0.05; Note S3) differential increases in transcript abundances (relative to t=0) for genes encoding photosynthesis-related proteins and significant differential decreases in transcripts encoding proteins involved in cell motility (Fig. 5A; SI Methods; Note S3). Among other highly up-regulated transcripts were genes associated with amino acid metabolism, cytochromes, iron-sulfur enzymes, and lipid metabolism, indicating significant restructuring of cellular metabolism and structure (Table S11). Many significantly differentially transcribed genes (77; 34% of genes with ļlog_2_(FC)ļ > 2) were present in the union of selected features. These patterns of differential transcript abundances were consistent with our hypothesis that mixotrophs such as prymnesiophytes can rapidly remodel their metabolism and shift their trophic strategy from encounter-driven phagotrophy to phototrophy in response to greater availability of limiting nutrients. We did not observe a similar shift in prymnesiophyte transcriptional patterns when nutrients were added to samples derived from the more nitrogen-rich, higher latitude waters of the transition zone (Table S12). Instead additions of iron or a combination of iron, nitrate, and phosphate to this community resulted in a restructuring of the photosynthetic machinery and included a significant decrease (*p_adj_* < 0.05) in transcripts encoding the iron-responsive protein flavodoxin (57) and a significant increase in transcripts encoding plastocyanin (58). Together, these results highlight the remarkable ability of mixotrophs to rapidly optimize their nutritional strategy to environmental conditions.

**Fig. 5.**
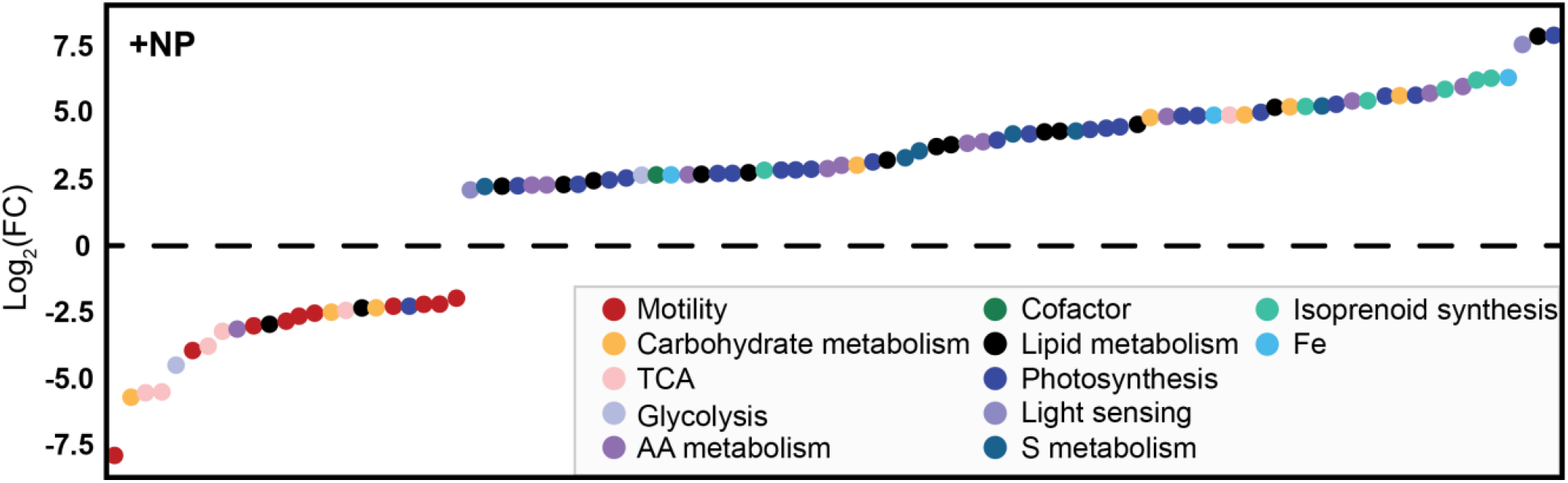
Transcriptional response of gyre community of mixotrophic prymnesiophytes to nitrogen and phosphorous amendment. Bottle incubations were performed on the Gradients 2 cruise (MGL1704, May 26 - June 13, 2017) with cells harvested for RNA extraction before incubation and 96 hrs after amendment with 0.5 μM PO_4_ and 5 μM NO_3_ (see SI Methods; Note S3). Transcripts with |Log_2_(FC)| > 2 and *p_adj_* < 0.05 are shown (224 total transcripts; Table S11). Transcripts are ordered by increasing Log_2_(FC) along the horizontal axis.

## Discussion

Our study introduces a machine learning driven approach that leverages transcriptional profiles to predict the *in situ* nutritional mode (heterotrophy, phototrophy, or mixotrophy) of protists in the natural environment. This method avoids potential artefacts associated with traditional, labor-intensive incubation-based estimates and opens the door for large-scale studies of how carbon flows through specific members of microbial communities. The model was trained with transcriptomes derived from protists grown under controlled laboratory conditions and challenged with a variety of validation transcriptomes. This approach identified a subset of gene family transcriptional profiles (features) that, given sufficient transcriptome replicates, resulted in trophic status predictions consistent with observed species-level nutritional strategies and broad phylogenetic patterns. The model accurately and consistently predicted the nutritional mode of heterotrophic or phototrophic specialists, whereas mixotrophy predictions were often coupled with those of either heterotrophy or phototrophy, likely reflecting both overlapping and distinctive mixotrophic attributes. The power of this approach is expected to increase with increasing diversity of transcriptomes available to train the model.

We predicted significant shifts in the trophic strategy of natural communities under different environmental conditions, both at a community level and for taxonomic bins corresponding to individual populations. Within the oligotrophic gyre, our model predicts that protists rely primarily on phagotrophy to acquire sufficient carbon and nitrogen for growth, a result consistent with optimal foraging theory, a resource allocation model (26), and a global ecosystem model that incorporates mixotrophy (30). Under nitrogen limitation, mixotrophs exhibit high intracellular C:N ratios and prioritize phagotrophy to acquire nitrogen from their prey (Fig. 5A). The lower intracellular C:N and C:P ratios of marine prokaryotes compared to protists (59) makes phagotrophy even better suited to fulfil the nutrient requirements of cells in the nitrogen-limited waters of the gyre. Given the presumed optimality of heterotrophic metabolism, why do mixotrophic organisms with both phagotrophic and photosynthetic capabilities dominate protist communities within the oligotrophic gyre? We propose that similar to what was observed in the amendment experiments, ephemeral injections of nitrogen into surface waters via mesoscale eddies, near-inertial waves, and internal tides (60, 61) rapidly shift the mixotrophic community to a more energetically favorable photosynthesis-based growth, thus preventing their displacement by heterotrophic specialists. During the dominant low-nutrient conditions, the ability of mixotrophs to consume prey (including photosynthetic prey) to fulfil their carbon and nitrogen requirements prevents their displacement by protist phototrophic specialists. The variable conditions of the gyre thus appear to mitigate potential costs associated with maintaining the complicated cellular machinery required for both photosynthesis and phagotrophy.

In the nutrient-rich North Pacific transition zone, a different scenario likely regulates the trophic status of the protist community. As nutrient concentrations increase, our model predicts that the community shifts from a primary reliance on phagotrophy in the gyre towards an increasing reliance on phototrophy and mixotrophy by smaller protists (0.2-3 μm) and mixotrophy by larger protists (3-200 μm). A similar shift with latitude is predicted for individual species, with increased predictions of phototrophy for the small chlorophyte *Micromonas* and mixotrophy for the larger *Chrysochromulina* species. The differing proportions of phototrophy predictions between size classes likely reflects the relationship between size-dependent uptake kinetics and nutrient availability, as smaller cells with a greater surface area:volume can more efficiently capitalize on lower dissolved nutrient concentrations (62). However, the metabolic theory of ecology predicts rates of heterotrophic metabolism will decrease disproportionately with decreasing temperature (63, 64). Why then do the larger mixotrophs continue to carry out phagotrophy in the cooler, nutrient-rich waters of the transition zone? Recent studies suggest that mixotrophic cells use the reductants generated via photosynthesis for organic carbon metabolism rather than carbon fixation (65). Moreover, the mixotrophs within our training set were distinguished by transcription of gene families encoding distinctive varieties of carbohydrate active enzymes. Thus, we propose that at the highest latitudes reached on the Gradients cruise, these larger mixotrophs fulfill their nutrient requirements via prey engulfment despite reduced sea surface temperatures, with necessary reductants for organic carbon metabolism supplemented through photosynthesis. We further propose that in accordance with results from the Edwards model (26), the reduced light levels we observed at the higher latitudes remain sufficient to have little impact on mixotroph abundance in surface waters.

Deciphering the functional role of mixotrophic protists in the marine carbon cycle has been a longstanding challenge, made difficult by the many caveats involved with current methods. Here we introduced a machine learning approach, made possible by the recent explosion of available transcriptomic data, and demonstrated that the model is skilled at inferring the trophic mode of natural populations based on the transcriptional patterns of select gene families. When we applied the model to metatranscriptomic data from the open ocean, predicted patterns in trophic status compare favorably with other results based on resource allocation models (26), global ecosystem models (30), the metabolic theory of ecology (66), and optimal foraging theory (55). By combining model predictions with bioinformatic analysis of metatranscriptomes obtained during on-board incubation experiments, we were able to develop intuitive explanations for observed functional differences between organisms and highlight the potential drivers of mixotroph ecosystem function. As the ubiquity of mixotrophy in the marine environment becomes increasingly apparent, so does the need to incorporate mixotrophs into our understanding of the ocean’s carbon cycle and the microbial ecology of the marine water column. Future coupling of our machine learning technique with targeted field experiments and numerical modelling will enable detailed dissection of the role that mixotrophs play in marine ecosystem processes.

## Methods

### Model training and evaluation

The MMETSP represents an imbalanced dataset for classification purposes (275 phototrophic, 93 mixotrophic, 78 heterotrophic, remainder unknown). We therefore carried out feature selection using four datasets consisting of randomly under-sampled phototrophic transcriptomes (*n*=80, 100, 120, 140) together with all mixotrophic and heterotrophic transcriptomes. Features that strongly impacted classification accuracy were determined for the Random Forest and XGBoost classifiers using the mean decrease accuracy (MDA) method (35). In MDA, the reduction in accuracy for a model is determined for each feature by randomly shuffling each feature across samples and carrying out 5-fold cross validation. In this approach, features that result in the largest mean decrease in accuracy are identified as key features. All features that resulted in a decrease in classification accuracy were retained. To examine clustered median expression of selected features in labelled MMETSP transcriptomes, data were reduced to the set of selected features, grouped by trophic mode, genes with zero median transcript abundances removed, and values were log-transformed. After feature selection, model performance was re-evaluated against the two reduced datasets (union and common set). The list of selected gene families (Pfams) is available in Table S2. Statistical measures were computed via the Kruskal-Wallis H-test with a post-hoc Pairwise Wilcoxon rank sum test, with the Benjamini-Hochberg correction for multiple comparisons.

### Validation transcriptome processing

Validation transcriptome read files (Table S6) were retrieved from NCBI’s short read archive through the SRA Toolkit and processed using an assembly and annotation pipeline similar to that presented in Johnson et al. (2018). Briefly, reads were quality controlled using Trimmomatic (v0.36) (67) and normalized using the normalize-by-median.py script from the khmer software package (68). Normalized reads were assembled with Trinity (v2.9.1) (69) and annotated via the dammit! pipeline (v1.2; https://github.com/dib-lab/dammit). Bowtie2 (70) was used to assess the percentage of quality-controlled paired-end reads that mapped back to the assemblies. For dammit! annotations, only matches to the Pfam database with an e-value < 10^-5^ were retained. Read counts in transcripts per million were generated for each assembly using Salmon (v1.2.1) in quasi-mapping mode.

### Metatranscriptomic data processing

Environmental transcriptome bins were obtained as follows: quality-controlled short reads were assembled using the Trinity *de novo* transcriptome assembler version 2.3.2 (69) on the Pittsburgh Supercomputing Center’s Bridges Large Memory system. Parameters included using *in-silico* normalization, a minimum k-mer coverage of 2, and a minimum contig length of 300. These raw assemblies were then quality controlled with Transrate v1.0.3 (71). Assemblies were merged and clustered at the 99% amino-acid identity threshold level with linclust in the MMseqs2 package to eliminate redundancy (72). Translated contigs were aligned to a reference sequence database of marine organisms including peptide sequences from hundreds of marine eukaryotic transcriptomes (73) using DIAMOND v 0.9.18 (74). Taxonomy was assigned with DIAMOND by using the top 10% of hits with e-value scores below 10^-5^ to establish the Lowest Common Ancestor of each contig. Putative function was assigned using hmmsearch (from HMMER 3.1b2 (75), using given trusted cutoff bitscores, --cut_tc) to find the best-scoring gene family from Pfam 31.0 (34). Contig abundances were quantified by pseudoalignment of the paired reads to the assemblies with kallisto (76) and normalized to the total assigned read pool of the taxonomic bin. Species-level transcriptional profiles were then normalized *in silico* to generate transcripts per million (TPM) profiles. Bin completeness was estimated by the number of non-zero transcript abundances within each bin. A completeness cut-off of 800 non-zero transcripts was selected based on the distribution present in the MMETSP dataset (see Note S1). Detailed information concerning RNA extraction and library preparation can be found in the SI Methods.

### Environmental metadata sourcing and processing

Environmental metadata and cruise data were obtained from the Simons Collaborative Marine Atlas Project pycmap API (CMAP; https://simonscmap.com/; Data originating from (52, 77–80)). Respiration rates were calculated following the equations presented in (81) using satellite-derived sea surface temperature (79) and photosynthetically available radiation (78) as input. Details of these rate calculations are presented in Supplementary Note 2. A detailed discussion concerning the origin and pre-processing of ancillary environmental metadata may be found in the SI Methods.

### At-sea incubation experiments

Twenty liters of seawater were collected into replicate polycarbonate carboys from 15m depth and incubated at *in situ* temperature for 96 hrs in on-deck, temperature-controlled incubators screened with 1/8-inch light blue acrylic panels to approximate *in situ* light levels at 15 m [55% of surface irradiance to approximate *in situ* light levels assuming an attenuation coefficient of 0.04 m^-1^ (82)]. Triplicate carboys were amended with 5 μM nitrate and 0.5 μM phosphate at *t* = 0. Duplicate carboys with no amendment served as a control. After 96 hrs, samples were filtered onto a 3 μm polycarbonate filter. Cells passing through the 3 μm filter were collected on a 0.2 μm polycarbonate filter. RNA extraction and sequencing were carried out as detailed in the SI Methods. Differential transcription analysis was performed in R using DESeq2 (83). Briefly, metatranscriptome reads from shipboard incubation experiments were mapped to prymnesiophyte reference transcriptomes using Salmon (84). The number of reads mapped from all treatments was consistently between 300,000 – 400,000. Differentially transcribed genes were identified using DESeq2 with an adjusted p-value cut-off of 0.05. A detailed discussion of this analysis is presented in Supplementary Note 3.

## Data availability

Scripts used for feature extraction, model training, and prediction are available at github.com/armbrustlab/trophic-mode-ml. Training data are available through Zenodo (10.5281/zenodo.4425690). Metatranscriptome data from the SCOPE Diel Cruise (KM1513) are available through the NCBI’s SRA under BioProject PRJNA492142, SCOPE Gradients 1 under BioProject PRJNA690573, and SCOPE Gradients 2 incubation experiments under BioProject PRJNA690575. *Chrysochromulina* sp. isolate transcriptomes are available under BioProject PRJNA690570 and PRJNA690572.

A detailed discussion of experimental and computational approaches is included in the SI Methods and SI Appendix.

## Acknowledgements

This work was funded by the Simons Foundation through Award #426570SP to E.V.A & A.E.W., XSEDE Grant allocation OCE160019 to R.D.G., and Simons Foundation Award #594111 to B.S.L. We thank G. Stewart and C. Schvarcz for providing *Chrysochromulina* sp. cultures used in this work, the SCOPE-Gradients team for their sample collection efforts at sea, K. Cain and F. Ribalet for flow cytometry analyses, N. Hawco and R. Bundy for assistance in the SCOPE Gradients 2 incubation studies, R. Morales and R. Lim for their expertise and effort carrying out RNA extractions, JB Raina for comments on an earlier version of the manuscript, and the captain and crew of the R/V Marcus G. Langseth and R/V Ka‘imikai-O-Kanaloa.

## Author contributions

B.S.L. and E.V.A. designed the research; B.S.L., R.D.G., M.J.S., S.C., B.P.D., A.J.A., and A.E.W. performed research. B.S.L, R.D.G, and M.J.S analyzed data. B.S.L. and E.V.A. wrote the paper with input from all co-authors.

